# Scalable platform for cellular and biochemical screening of combinatorial small-molecule libraries in droplets

**DOI:** 10.64898/2026.06.12.730915

**Authors:** Mihajlo Todorovic, Laura West, Frederic Daniel Sigoillot, Gianluca Etienne, Asitha Adrian Abeywardane, Antonin Tutter, Tushar Apsunde, Douglas S. Auld, Andrew Brady, Seth Carbonneau, Lia Ecker, Jian Fang, Christophe Freslon, Jacob Hale, Hidetomo Imase, Fupeng Ma, Brian Minie, Steven Paula, Andrea K. Pomerantz, Piro Siuti, Charles A. Wartchow, Ken Yamada

## Abstract

A new pattern of hit discovery is emerging with the rise of chemocentric modalities, such as chemical induced proximity (CIP). Synthesis and functional screening of dynamic, purpose-built libraries, such as E3 ligase biased libraries for targeted protein degradation (TPD), is redefining the druggable landscape. However, this new paradigm arguably lies beyond the remit of conventional screening methods which have been optimized for static, diversity-oriented libraries. To address this gap, we developed microfluidic compound screening in droplets (MicDrop), a scalable platform where purpose-built DNA-encoded one-bead one-compound library members are individually tested at high concentrations in pico-liter sized droplets for either biochemical or cellular function. A proof-of-concept CRBN library demonstrated the robustness of the workflow through an IKZF3 degradation screen which discovered an unexpected succinimide-based IKZF3 degrader, while reproducing the relative potency ranking of reference compounds. Screening of a prospective VHL library for CDO1 recruitment and degradation demonstrated that orthogonal assays gave an informative consensus hit list and an actionable machine-learning (ML) recruitment model. These findings establish a blueprint for intentional discovery of chemical inducers of proximity, while producing reliable data to accelerate ML-based design-make-test-analyze (DMTA) cycles in pursuit of new therapeutics.

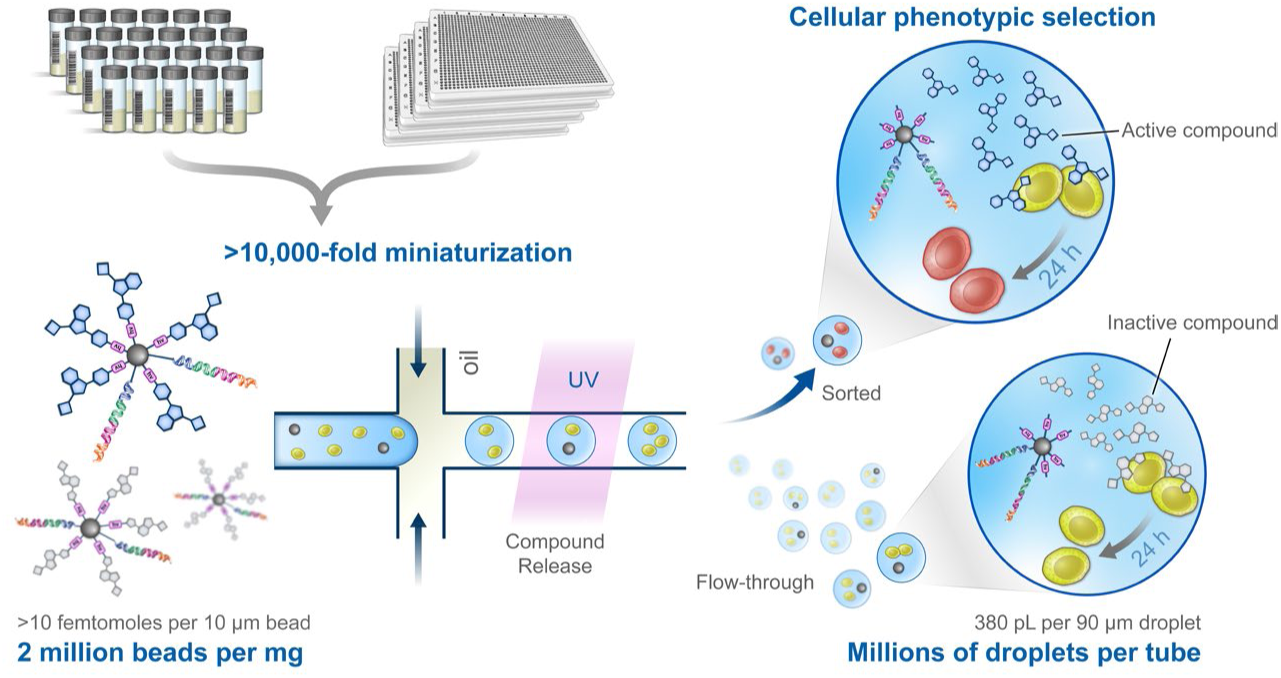

## Introduction

Empirical screening of large compound collections remains a cornerstone of modern drug discovery. Recent advances in machine learning (ML) models only accentuate the importance of large, curated, and relevant data generated for such endeavors^1–3^. Concurrently, the rise of molecular glues that expand druggability via chemical induced proximity (CIP)^4,5^ with *intentional* design of molecules^6–8^ has revealed the acute need for scalable synthesis of novel bespoke libraries, biased for a particular effector or protein of interest, and their cellular functional screens to monitor implicated phenotypes. Such a paradigm could deliver advanced, progressable starting points for rapid optimization towards clinical candidates, preferably establishing direct-from-screen structure-activity-relationships (SAR) and accompanying machine-learning (ML) activity models that bypass laborious hit resynthesis and validation^2,3^.

Molecular glue degraders (MGD) that elicit targeted protein degradation (TPD) represent a prototypical CIP modality that is redefining what is druggable by small molecules^5,6,8–17^. For example, Novartis has strategically invested in building large cereblon (CRBN)-biased libraries numbering in the thousands to tens of thousands and identified molecular glue degraders for a variety of previously undruggable targets^18–20^, an approach also reported by others^7,10,21–25^. To the best of our knowledge, these compounds were synthesized, purified and characterized one at a time before being tested in microtiter well plate format in traditional high-throughput screens (HTS), a highly resource intensive endeavor. To address the synthesis and purification bottleneck, various high-throughput chemistry approaches have been developed, most notably purification-free, direct-to-biology (D2B) methodologies yielding promising early results^26–30^. However, D2B approaches typically employ single-step chemistry with diversity and required effort that scales linearly, typically only to thousands, leaving a significant gap to reach the diversity and scale required for intentionally discovering molecular glues for a diverse set of targets.

In recent decades, synthesis and screening of DNA-encoded libraries (DELs)^31–38^ has become a mainstream small-molecule ligand discovery method^38–40^ granting access to vast combinatorial chemical space in the millions to billions via split-pool chemistry while requiring minimal instrumentation and compound management^38^. DELs enable massively pooled screens through the covalent linkage of the combinatorially synthesized small molecules to the DNA tags. However, this covalent tagging also limits their application to affinity selections with immobilized biological targets, requiring resynthesis off-DNA for validation of their true activity in cellular and biochemical milieu. Although tailored DEL selection methods have been devised to select for recruitment^41,42^ and ubiquitination^43,44^ for TPD applications, as well as for GPCR agonism^45^, a direct and scalable platform for cellular functional or phenotypic selection of DELs remains elusive.

To combine the direct biological readout and replicate-based robustness offered by traditional microtiter plate-based screens with the chemical enumeration power and scalability of DELs, we sought a miniaturized platform with discrete compartments capable of hosting a combinatorially prepared library member, which can be released into a confined solution, and reporter cells to read out a desired phenotype. Among several possible systems such as agar lawns (500∼1,900 beads per dish)^46,47^, nanopens (3,500∼20,000 pens per chip)^48,49^ picowells (∼88,000 wells per chip)^15,50^ or microcapillaries (∼800,000 capillaries per chip)^51,52^, we opted for water-in-oil microdroplets (millions in a tube)^53,54^, for which operational simplicity and scalability stood out.

We were particularly inspired by the pioneering work of Paegel and colleagues^55^ where each DNA-encoded one-bead one-compound (eOBOC) library member releases many copies of a single, combinatorially synthesized molecule within a droplet compartment upon photoirradiation, thereby performing discrete compound screening in an isolated compartment, directly analogous to an individual well in traditional HTS. Millions of microcompartments can be pooled in a single tube and sorted continuously on the basis of a biochemical readout in flow using fluorescence activated droplet sorting (FADS). The system has been successfully employed to screen for inhibitors of recombinant enzymes^56–58^ and *in-vitro* transcription translation (IVTT) systems^59^. However, extrapolation of this droplet screening platform to cellular systems has remained elusive, due to compounding challenges, including; compound retention^60,61^, assay relevance^62^, library bead clumping^55^ and auto-fluorescence^57^, low fraction of droplets with one cell and one bead due to double-Poisson statistics^55,63,64^, and inherent variability of fluorescence reporter expression and response levels^64^. Our solutions to these challenges will be detailed in due course.

In this manuscript, we describe a general platform for microfluidic compound screening in droplets (MicDrop, Figure 1) and demonstrate its application to two independent proof-of-concept (PoC) molecular glue systems for TPD, namely, IMiD-induced recruitment and degradation of IKZF3 via CRBN^9,65^, and VH-032-induced recruitment and degradation of CDO1 via VHL^16^. First, the robustness of the workflow was demonstrated with a replicate rich PoC CRBN-biased library screened for cellular IKZF3 degradation, where the enrichment directly correlated with relative potency in an orthogonal plate-based assay, while also identifying a novel succinimide-based IKZF3 degrader. In addition, a prospective VHL-biased library was screened for cellular CDO1 degradation and biochemical CDO1-VHL recruitment. The pair of screens gave a consensus hitlist with a high hit confirmation rate and a machine learning model trained and benchmarked only on droplet screen data demonstrating robust performance by delivering actionable structure-activity relationship (SAR).

**Figure 1:**
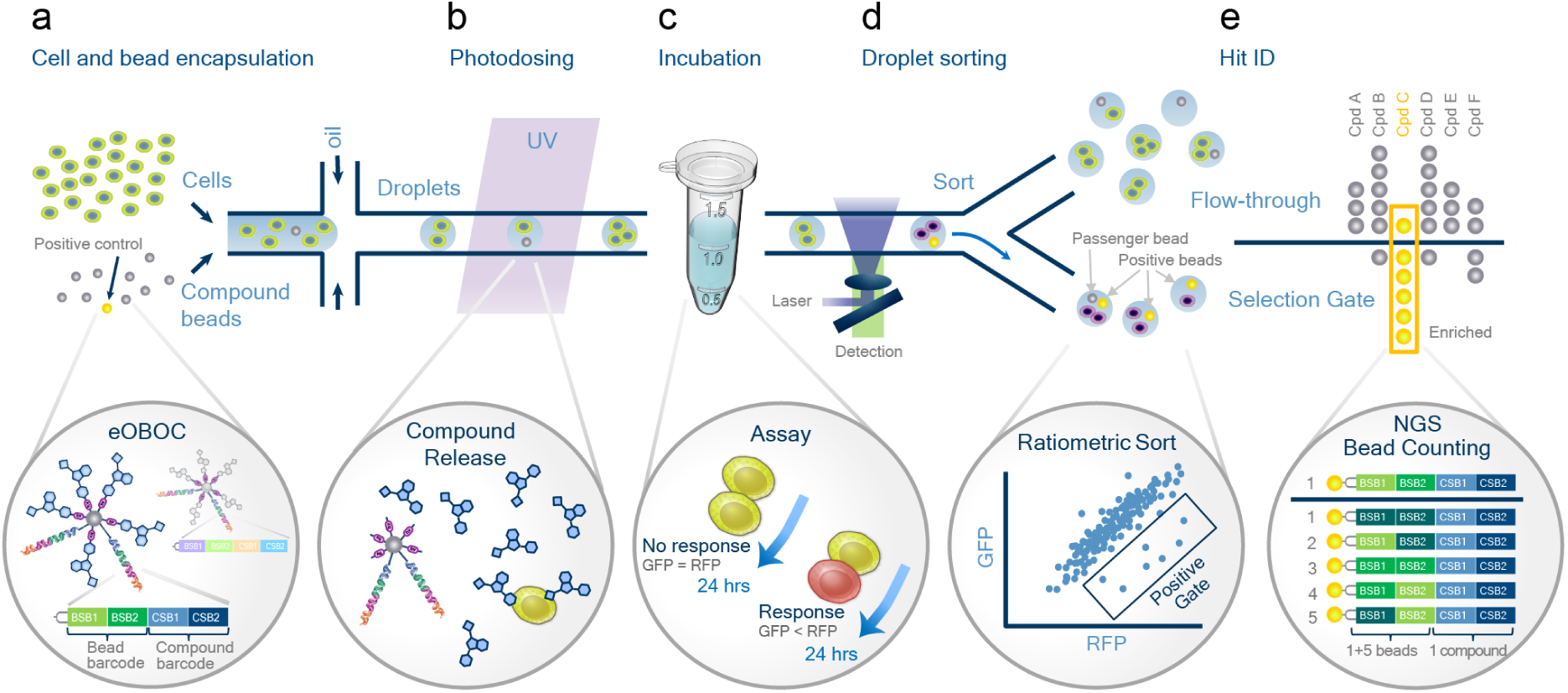
Schematic representation of a MicDrop workflow: Phenotypic screen of combinatorial library released from DNA-encoded carrier beads in droplets. From left to right; a) bead and cell encapsulation; a premade suspension of beads and cells is flowed into a flow-focusing microfluidic chip at a continuous rate, meeting a stream of fluorinated oil at the cross-junction, resulting in the production of monodisperse oil-in-water droplets, a plurality of which contain at least one cell and one or fewer beads. Each bead carries many copies of the same compound and DNA tag encoding for bead identity (BSB) and compound identity (CSB). b) The droplets then flow into a UV dosing chamber, where 365 nm light releases a tagless compound from the photocleavable linker loaded on the bead. c) The droplets are then collected into an Eppendorf tube and stored at the appropriate temperature and humidity for the required duration of the assay. d) The droplets are then sorted by fluorescence activated droplet sorting (FADS), based on a ratiometric readout between green fluorescence (488 nm excitation, 543 nm +/− 11 nm emission filter) and red fluorescence (561 nm excitation, 591.5 nm +/− 21.5 nm emission filter), where the droplets with lowest green/red ratio are collected in the selection gate and the remaining droplets in flow-through. e) The beads are isolated from the collected droplets from the selection gate and the flow-through and the DNA barcodes amplified and sequenced by NGS. Bead specific barcodes allow for accurate bead count estimates per compound in selection gate vs. flow-through, enabling determination of bead enrichment and statistical analysis.

## Results

### IKZF3 ZF2 β-hairpin-GFP/RFP sensor assay in droplets

For an IKZF3 degradation assay compatible with a fluorescence-activated droplet sorter (FADS), we selected a GFP/RFP ratiometric assay where GFP is fused to IKZF3 zinc finger 2 (β-hairpin), or more simply, IKZF3-GFP/RFP sensor assay (Figure 2a). IKZF3 zinc finger 2 is a well-characterized degron^9^, which is sufficient to elicit degradation of its fusion partner protein upon IMiD drug treatment^66^. Because of its ability to normalize for varying expression levels of the protein of interest, this sensor assay format is often utilized for studying cellular protein degradation with fluorescence activated cell sorters (FACS)^9,67^. In our case, a construct featuring an IKZF3-AcGFP fusion and DsRed in the same open-reading frame, separated by a self-cleaving P2A site, was expressed in Jurkat cells. The shared promoter ensured that cells exhibit a constant ratio of green and red fluorescence at baseline. In the presence of a glue degrader such as Compound 1, the level of IKZF3-GFP fusion protein would decrease, resulting in a loss of green, while maintaining red fluorescence signal (Figure 2a).

**Figure 2:**
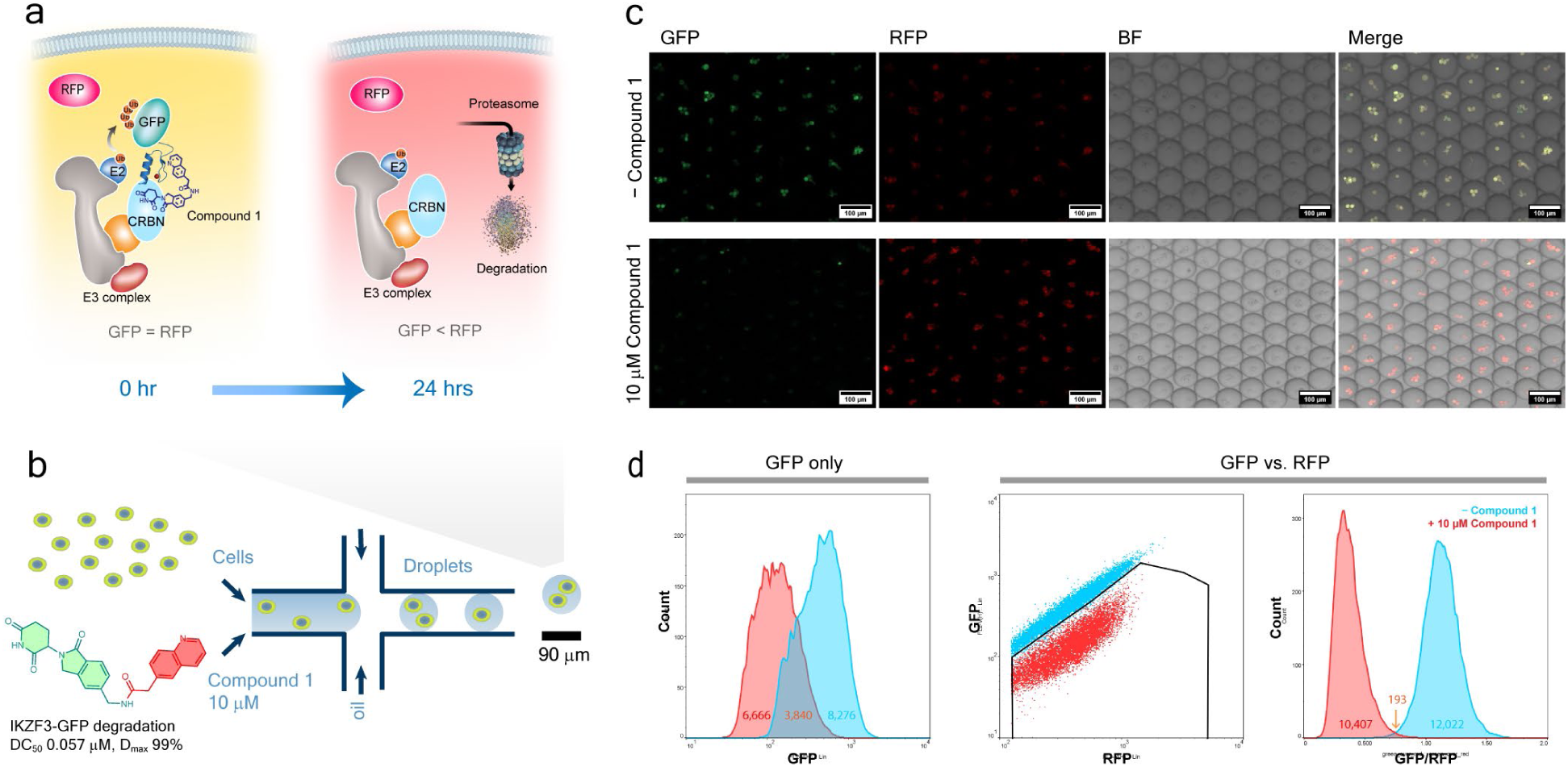
Validation of IKZF3-GFP/RFP sensor assay in droplets. a) Schematic representation of ratiometric IKZF3-GFP/RFP sensor assay. Compound 1, a molecular glue degrader of IKZF3 (DC_50_ 57 nM, D_max_ 99%), binds to CRBN, part of the E3 ligase complex, and recruits the IKZF3 β-hairpin, promoting the poly-ubiquitination of the fusion protein and its subsequent degradation by the proteasome resulting in loss of green, while maintainging red fluorescence that acts as an internal standard. b) Schematic representation of cell and reference compound co-encapsulation into water-in-oil droplets using a flow focusing microfluidic device. Every droplet contains 10 µM Compound 1 and, on average, 2.3 cells are loaded into a single droplet. c) Assay readout in droplets by confocal microscopy 24 h after generation, from left; green channel (GFP), red channel (RFP), brightfield (BF) and overlay (merge). From top; without and with 10 μM Compound 1. d) FADS plot from On-chip Sort (On-chip Biotechnologies). Left: Overlay of FADS histograms of IKZF3-GFP/RFP sensor assay 24 h after droplet generation, droplet count vs. green fluorescence, with (red) and without (blue) Compound 1 treatment. Middle: Overlay of FADS scatter plots of green vs. red fluorecence, with (red) and without (blue) Compound 1 treatment. Right: Overlay of ratiometric FADS histograms of droplet count vs. green/red ratio (threshold ratio 0.75), with (red) and without (blue) Compound 1 treatment.

To validate the IKZF3-GFP/RFP sensor assay in droplets, a PDMS-based flow-focusing microfluidic chip was utilized to encapsulate cells with compound loaded assay media in monodisperse droplets with a diameter of 90 µm (Figure 2b). For the continuous phase, we used a commercially available, bio-compatible fluorinated oil and surfactant (2% 008-FluoroSurfactant in HFE7500, RAN biotechnologies). The generated droplets were incubated at 37 °C for 24 hours in an Eppendorf tube before sorting by a commercial FADS (On-chip Sort, On-chip Biotechnologies), which was used for all FADS experiments described in this manuscript. The encapsulated Jurkat cells expressing the IKZF3-GFP/RFP sensor assay responded as expected to the presence of 10 µM of a potent IKZF3 degrader, Compound 1 (DC50 57 nM, Dmax 99%), with a loss of green fluorescence after 24 hrs incubation, while maintaining red fluorescence, as observed by both microscopy (Figure 2c) and FADS (Figure 2d). Five additional IKZF3 degraders with varying potency and a negative control were also tested and showed IKZF3-GFP degradation (data not shown) roughly correlating with their respective maximal percentage degradation (D_max_) determined by standard plate-based assays (Figure 3c).

**Figure 3:**
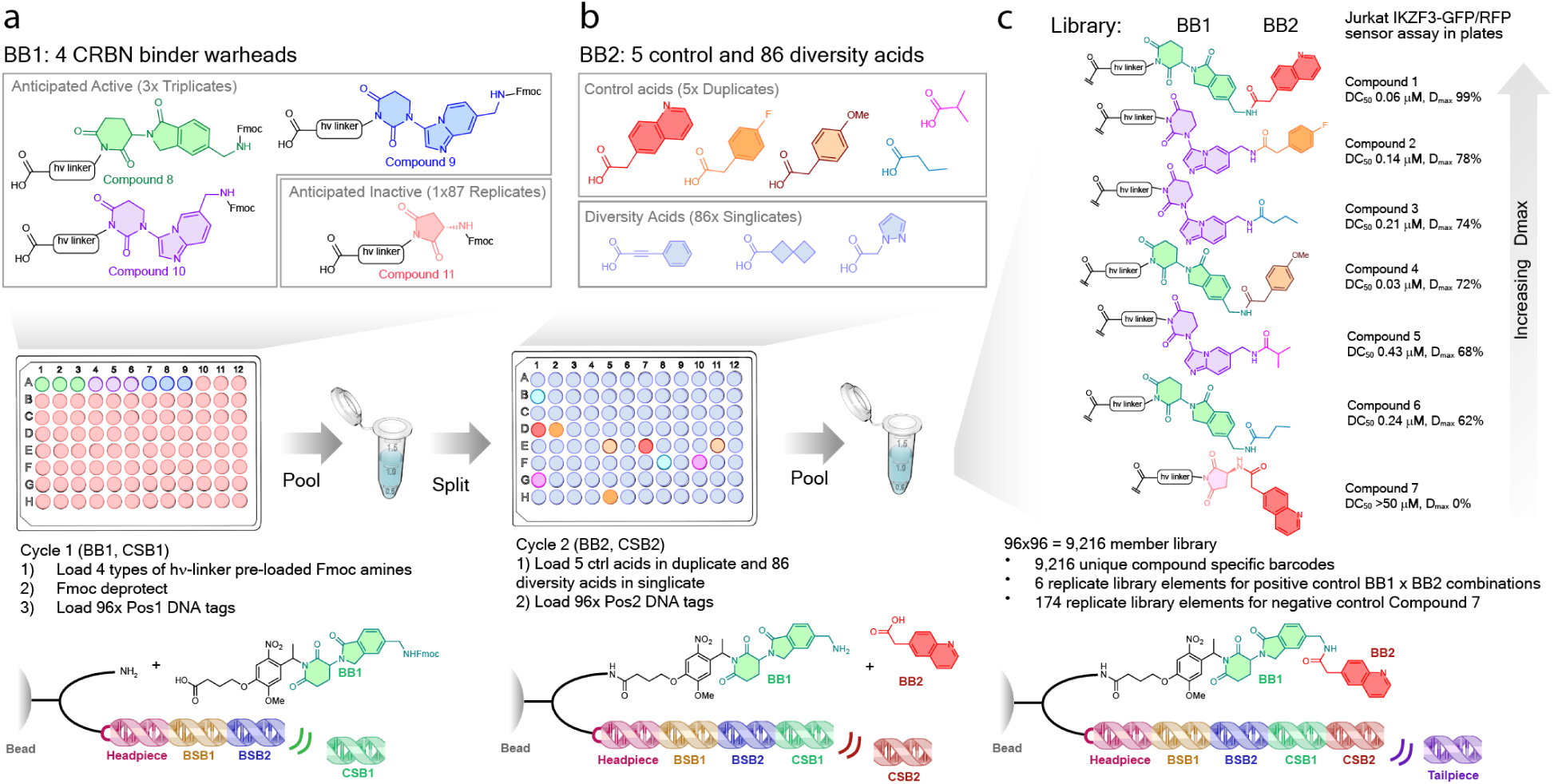
Design and synthesis of the 9,216-element CRBN-biased PoC library. a) The first building block (BB1) position contained four known CRBN binder motifs Fmoc-protected and preloaded on the photocleavable linker, three of which were validated IKZF3 degrader motifs (green, purple and blue) each represented thrice and an amino-succinimide not expected produce IKZF3 degraders (tan) represented 87 times. The acids were first loaded onto pre-barcoded (BSB1xBSB2) beads in each of the 96-wells, followed by Fmoc deprotection, BSB1 DNA-tag (CSB1) ligation and pooling. b) The second building block (BB2) position contained the 5 control carboxylic acids (red, orange, brown, blue and cyan) each in duplicate, as well as 91 diversity-oriented carboxylic acids (examples in light blue) in singlicate, with each replicate coded for by one of 96 unique barcodes (CSB2). The pooled beads from the previous step were split into 96 wells, and the acids were loaded in each of the 96 wells, followed by CSB2 DNA-tag ligation and pooled. c) The resulting library was ligated with tailpiece to give a 9,216 element, 364-member library with 9,216 unique CSB1 x CSB2 barcode combinations. The six positive controls Compounds 1-6 are represented 6 times each, and the negative control Compound 7 is represented 174 times each (shown). The remaining 357 combinations are not shown.

The GFP/RFP sensor assay addressed three key challenges in developing a cellular functional assay in droplets. First, it accounted for signal variation at both the cell-to-cell and droplet-to-droplet level, owing to the equal ratio of GFP/RFP at baseline and ratiometric readout, respectively. This improved resolution is evident when comparing green fluorescence alone vs. green-red fluorescence ratio (Figure 2d, left vs. middle/right). In addition, no longer bound to targeting for a single cell per droplet, we could substantially increase the cell loading, thereby ensuring approximately 90% of droplets had at least one cell. This, in essence, reduces the double-Poisson problem of previously envisioned cellular eOBOC screens^55,64^ to a single-Poisson distribution, increasing the sorting efficiency and achievable throughput by approximately 5-fold, while also substantially reducing false-negatives arising from bead-only droplets.

### Proof-of-concept 9,216-element CRBN-biased glue library with 384 unique members

The library bead chemistry was adopted from previously described eOBOC systems^55,56,68^ with some modifications (Figure 3, bottom). The tag starts with a headpiece containing the forward primer followed by a bead specific barcode (BSB), which is a combination of two modules: BSB1 and BSB2, prepared via split-pool ligation prior to compound loading. The compound specific barcode (CSB) follows, which in turn consists of a combination of two modules: CSB1 (encoding BB1), which was ligated on following the incorporation of the first building block (BB1), a CRBN binding motif pre-loaded on a nitroveratryl linker, and CSB2 (encoding BB2) which was ligated on following the incorporation of the second building block (BB2), a carboxylic acid capping motif. Each unique combination of CSB is referred to as a library element throughout this manuscript. The DNA tag is then capped with a tailpiece that includes unique molecular identifier (UMI) and reverse primer to complete the barcode.

The PoC library was designed to include building block redundancy at multiple levels, particularly emphasizing the six positive controls (Compounds 1-6) and the negative control Compound 7 (Figure 3a-c). This allowed us to compare the performance of individual compounds coded for by multiple redundant barcodes or aggregate the results for added statistical power. Thus, for the first cycle comprised of 96 elements, 4 CRBN binder warheads were selected, with glutaramide (Compound 8) or dihydrouracil (DHU) (Compound 9 and Compound 10) warheads represented in triplicate and succinimide (Compound 11) warhead representing the remaining 87 positions (Figure 3a). Based on the prior structure-activity relationship (SAR) known at the time of library design, Compound 8 and Compound 10 contained motifs present in known potent IKZF3 degraders, Compound 9 was only known from weak IKZF3 degraders and Compound 11 was not expected to produce any active combinations.

For the second cycle comprised of 96 elements, the five acids contained in the previously known positive controls (Compounds 1-6) were represented in duplicate, with diversity acids representing the remaining 86 elements in singlicate (Figure 3b). The result was a 9,216-element (BSBs) encoded library comprised of 364 unique compounds, including each of the 6 positive controls (Compounds 1-6) represented by 6 replicate library elements, and negative control Compound 7 represented by 174 replicate library elements, with an expected hit rate of 1-2% based on our prior SAR knowledge (Figure 3c).

The library was synthesized by a modified protocol based on previously reported split-pool methodology in 96 well plates^68,69^, with approximately 300 million beads total. To ensure the overall quality of the encoding DNA tags, qPCR was performed on the library post synthesis, finding an approximate loading of approximately 18 million tags per bead. Additionally, preliminary NGS analysis confirmed all 96 possible barcodes for each of the BSBs (code 2-3) and CSBs (codes 4-5) were represented.

### Cellular screen of 9,216-element PoC CRBN-biased glue library for IKZF3-GFP degradation

With the library in hand, we performed a cellular IKZF3 degradation screen with 3 independent runs using the GFP/RFP sensor assay. A solution consisting of library beads and cells in aqueous media was encapsulated into droplets (∼90 µm) using a microfluidic device and treated, in-line, with 365 nm UV light to liberate compounds into solution within each droplet, resulting in free-ligand concentrations above 10 µM. Overnight incubation was then followed by droplet sorting based on GFP/RFP ratio (Figure 4a). Input for each run was approximately 1,000,000 beads or 100 library equivalents. Following sorting, approximately 15,000 beads from the selection gate (top 5%) and approximately 400,000 beads from the flow-through were isolated (Figure 4b). All the beads from the selection gate and a sample from each corresponding flow-through, approximately 100,000 beads each (Figure 4b), were analyzed for hit calling.

**Figure 4:**
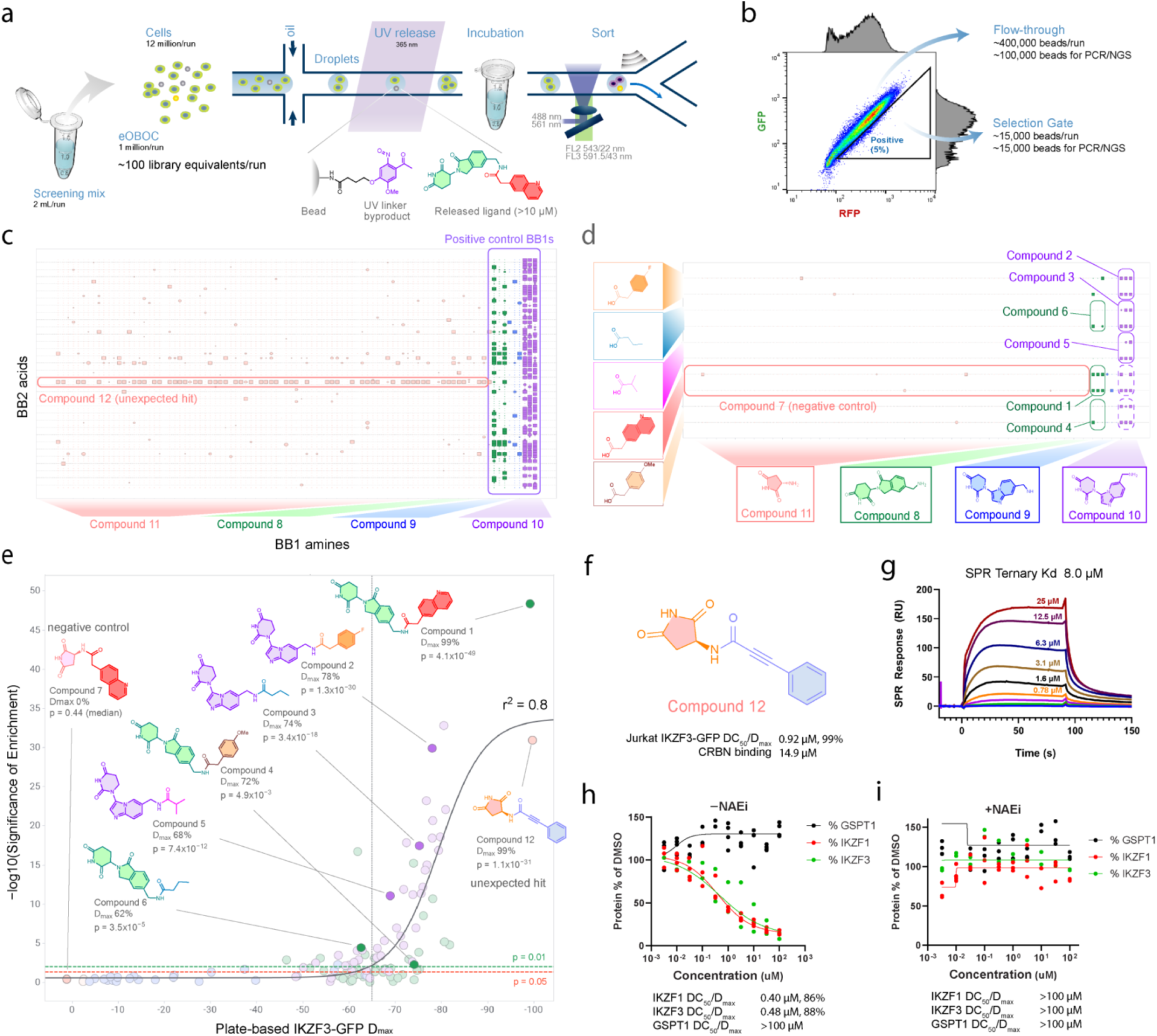
Cellular screen of 9,216-element PoC CRBN-biased glue library for IKZF3 degradation. a) Schematic for cellular IKZF3 degradation screen in droplets. A screening pre-mix of library beads and cells is encapsulated in droplets using a microfluidic device. Inline UV dosing activates the photocleavable linker and releases free ligands within the droplet compartments (>10 µM compound concentration). The droplets are then incubated overnight before they are sorted by FADS based on GFP/RFP ratio. All the beads in the selection gate and a fraction of beads in the flow-through are subjected to PCR/NGS and data analyzed for hit calling. b) FADS plot of a typical sorting experiment. Approximately 5% of all qualified droplet populations are sorted into the selection gate. c) 2D plot of building block enrichment by statistical significance. Amine building blocks (BB1) are plotted on the x-axis and acid building blocks (BB2) are plotted on the y-axis. Combined result of run 1 and run 2. Fisher’s exact test (square; p < 0.01, circle; 0.01 ≤ p < 0.05, line; p ≥ 0.05). d) close-up of Figure 4b focusing on the individual CSB replicates of 4 BB1 amine x 5 BB2 acid combinations, including all 6 positive controls and the negative control. e) Correlation plot with independently synthesized compounds tested in plate-based high-content imaging version of the same IKZF3 degradation assay. Statistical significance of hit enrichment vs. maximal percent target protein degradation (D_max_) in well-based assay. CSB replicates are aggregated. r^2^ = 0.80. One compound was excluded from correlation based on presence of interfering particulate in the plate-based IKZF3 degradation assay. f-i) Characterization of Compound 12: g) SPR sensorgram of immobilized CRBN-DDB1 complex with constant flow of 20 µM Compound 12 and titration of GFP-Hild fusion protein. h-i) Capillary Western dose-response in OPM2 cells with antibody staining for IKZF1, IKZF3 and GSPT1, h) in absence and i) in presence of Neddylation inhibitor (NAEi) MLN4924.

The DNA barcodes on the beads were amplified by PCR, purified, and then submitted for paired end Illumina NGS sequencing. The sequencing read pairs were quality-filtered and merged, further filtered for expected barcode length and PCR duplicates were deduplicated, taking advantage of a UMI presence in the pre-processed sequences The pre-processed sequences were then aligned separately to expected bead specific barcodes (BSB: BSB1 x BSB2 combination) and to expected compound specific barcodes (CSB: CSB1 x CSB2 combination). Alignment results were merged so each preprocessed sequence was assigned a corresponding BSB x CSB combination. The data was aggregated as the number of preprocessed reads (UMI count) for each BSB x CSB combination and the combinations were sorted from the highest UMI count. The BSB x CSB combinations were given an index number starting with 1.

To define the discrete number of beads (BSBs) observed by NGS per CSB, which we refer to as bead count, we looked for the number of unique combinations of bead barcodes (BSB: BSB1 x BSB2 combination) per given library element (CSB1 x CSB2) which were found in the top indexed beads up to the number of expected bead count per sample as determined by a Countess cell counter. With this method, we detected 8,735 of the expected 9,216 library elements (95% coverage), with a median UMI count of 204 (when all the sequenced beads across all samples from the screen were combined). For the hit calling, we calculated the relative abundance (bead count) of a given library element in the positive-gate vs flow-through, based on the expected bead count and performed a Fisher’s Exact test^70^ to determine the statistical significance (p-value) of enrichment.

### Hit calling for 9,216-element PoC CRBN-biased glue library for IKZF3-GFP degradation

Among several analyses that were performed, a 2D plot of unaggregated BB1 amines (CSB1) vs. BB2 acids (CSB2), where the presence of a square signified that the given compound was significantly enriched (p value <0.05), provided an intuitive view of the overall hit landscape and replicate reproducibility (Figure 4c). This plot is analogous to a frequently employed method of hit analysis for the conventional affinity selection DEL paradigm^38^. The overall hit rate of our screen was 4.2%, or 367 out of 8,694 detected individual library elements, with a stringent hit threshold of p<0.01. This hit rate was higher than anticipated from the library design (1-2%) potentially because 1) we assumed D_max_ >80% would be required for a strong hit whereas we see many hits with p<0.01 with lower D_max_, and 2) there were many more active BB2 acid combinations with known active BB1s than was expected based on our previous SAR knowledge.

Next, we examined the BB1 level enrichment (Figure 4c, vertical lines). Treating individual library elements as a unique compound without aggregation, 6 of the 96 BB1s stood out as having many active combinations (the 3 library element replicates of Compound 8 and Compound 10). This result aligned well with our expectations given the library design and SAR knowledge. Each of the 87 replicates of intentionally inactive amine (Compound 11, tan) showed a low frequency of hits, matching the approximate rate expected from random events when the p value threshold is set at 0.01 (86/7,943 or 1.1%). The amine building block with only weakly active combinations known, (Compound 9, blue), showed a low frequency of hits slightly above the noise level (8/216 or 3.7%), whereas the amine building blocks with many known active combinations (Compound 8, purple, and Compound 10, green) showed a very high frequency of hits with a wide variety of BB2 acids (Compound 8; 45/258 or 18.2%, Compound 10; 148/288 or 51.4%). Thus, at the BB1 level, the general performance of the end-to-end workflow was confirmed to be very robust.

Although the succinimide BB1 (Compound 11, tan) was not known to yield any IKZF3 degraders, its combination with BB2 phenylpropiolic acid stood out, displaying a horizontal line (Figure 4c), with 56/83 replicate library elements (67%) showing highly significant enrichment (p<0.01). We therefore decided to follow up on this potentially unprecedented succinimide based glue degrader, Compound 12, for further validation (see “characterization of new hit, Compound 12”).

We next sought to understand the reproducibility of the individual replicates of the reference compounds (Figure 4d). The most potent positive controls Compound 1 and Compound 2 showed significant enrichment in 6 out of 6 replicate library elements, with 100% reproducibility across the replicates. On the other hand, the moderate and weak positive controls showed varying frequencies of significant enrichment, ranging from 1/6 (17%) for Compound 4 to 5/6 (83%) for Compound 3. For the negative control, Compound 7, we observed significant enrichment in only 2/166 replicate library elements (1.2%) consistent with the hit selection threshold p <0.01 (accepts 1% false positive). Thus, while the replicate library element reproducibility is robust for the most potent and inactive degraders, it becomes more variable for weaker degraders.

To understand the relationship between the number of beads screened per compound and the statistical power for hit calling, we leveraged the library design with intentional library element (synthesis) replicates that could be aggregated for analysis. Thus, aggregation of 6 replicates of each of the six positive controls (Compounds 1-6) resulted in highly significant enrichment (p <0.01) in all instances, ranging from 4.1×10^-49^ (Compound 1) to 0.0049 (Compound 4), whereas aggregating 166 replicates of the negative control (Compound 7) similarly to the positive controls did not give significant enrichment (median p = 0.44) (Figure 4e).

### Benchmarking cellular IKZF3 degradation screen of 9,216-element proof-of-concept CRBN-biased glue library

To understand the broader performance of our droplet-based cellular screen workflow, we characterized 107 independently synthesized and purified compounds featured in our PoC library using a plate-based high-content imaging version of the same IKZF3 degradation assay in Jurkat cells to determine their D_max_ and DC_50_. The significance of enrichment strongly correlated with plate-based IKZF3 degradation D_max_ at both the individual library element level (data not shown) and more robustly after aggregating library element replicates (r^2^ = 0.80, Figure 4e). The described workflow is thus able to semi-quantitatively rank order the hits with statistics derived from bead replicates.

We also determined the precision (#true positives/{#true positive + #false positives}) and recall (#true positives/{#true positives + #false negatives}) for the aggregated library element case, with the droplet screen threshold of statistical significance of enrichment set at a p-value of 0.01 and with a plate-based assay threshold D_max_ of 65% to qualify as a true positive in the confusion matrix (Figure 4e). In this case, we identified 43 selected positives of the 57 actual positives, resulting in a recall of 0.75 (57/76). We also found 14 false positives, also resulting in a precision of 0.75 (43/54, data not shown).

### Characterization of new hit, Compound 12

Next, we set out to validate the unexpected succinimide-based hit, Compound 12 (Figures 5f-i). First, we confirmed potent degradation of IKZF3 by Compound 12 in the Jurkat GFP/RFP sensor assay (DC_50_ = 0.92 μM, D_max_ = 99%), concurring with the observed enrichment. Next, we confirmed weak but dose-dependent CRBN binding of Compound 12 in a lenalidomide-based probe displacement FP assay^18^ (14.9 μM). To validate selective, on-mechanism degradation, we next performed capillary western with Compound 12 for degradation of native proteins, where Compound 12 selectively degraded IKZF1 and IKZF3 (DC_50_ = ∼0.5 μM for both), sparing GSPT1 (DC_50_ >100 μM) (Figure 4h). Degradation of IKZF1 and IKZF3 was rescued with neddylation inhibitor (NAEi) MLN4924 (Figure 4i).

**Figure 5:**
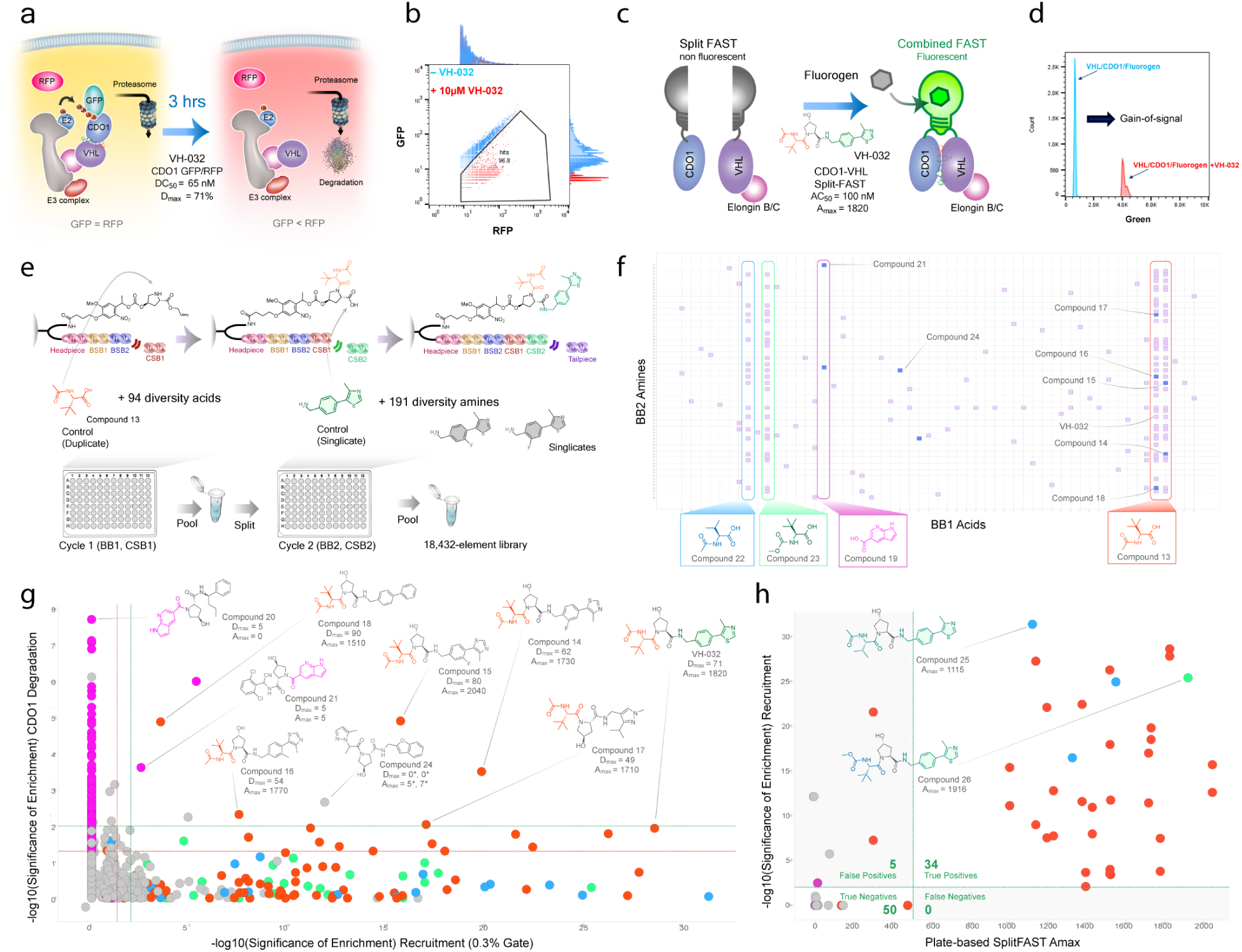
VHL library design, screen and hit validation. a) Schematic representation of cellular CDO1-GFP/RFP sensor assay: Analogously to the IKZF3-GFP/RFP sensor assay, the CDO1 glue degrader VH-032 (DC_50_ = 0.065 μM, D_max_ = 71%) recruits CDO1 to the VHL-E3 ligase complex, leading to poly-ubiquitination and proteasomal degradation of CDO1-GFP and subsequent loss of green fluorescence, but not red fluorescence. b) FADS plot of droplets containing CDO1-GFP/RFP cells post 3 hour incubation at 37 °C with (red) and without (blue) 20 μM VH-032. ∼20% of droplets were bright enough to register. Green channel: 488 nm excitation, 543 nm +/− 11 nm emission filter. Red channel: 561 nm excitation, 591.5 nm +/− 21.5 nm emission filter. c) Schematic representation of the splitFAST biochemical assay: In the presence of a CDO1-VHL glue such as VH-032 (AC_50_ = 100 nM, A_max_ = 1820), the two fragments of the splitFAST protein, each fused to either CDO1 or VHL, are brought in close proximity to form FAST protein which in turn activates the fluorogen to become highly fluorescent upon binding. d) FADS plot showing comparison of signal intensity for 10 μM VH-032 and DMSO control (488 nm excitation, 543 nm +/− 11 nm emission filter). e) VHL library design: The first BB was one of 95 diversity acids in singligate or one control acid from VH-032, acetyl *tert*-leucine, in duplicate. The second BB was one of 192 diversity amines in singlicate. Among the amines, motifs known to degrade CDO1 such as the (4-(4-methylthiazol-5-yl)phenyl)methanamine in VH-032 were included. They were loaded on BSB barcoded beads pre-loaded via carbamate to dual-protected hydroxyproline. Sequential deprotection of Fmoc and BB1 loading and CSB1 ligation, pool and split, followed by allyl ester deprotection, BB2 amine loading and ligation of CSB2, pooling and ligation of closing sequence completed the library. f) 2D enrichment plot for the splitFAST biochemical recruitment screen, with BB1 acids plotted on the x-axis and BB2 amines plotted on the y-axis. Only compounds with a p-value of 0.01 or below for the biochemical recruitment screen are represented with squares. Purple shading represents significance of enrichment (p value <0.01) in splitFAST recruitment screen (0.3%) gate, blue shading represents significance of enrichment (p value <0.01) in both splitFAST recruitment screen (0.3%) and the combined CDO1 cellular degradation screen. g) 2D pivot plot: Significance of enrichment in the CDO1 cellular degradation screen (all 4 screen runs combined) is plotted on the y-axis. Significance of enrichment in the splitFAST biochemical recruitment screen (only the 0.3% selection gate) is plotted on the x-axis. The green dashed lines represent a p-value threshold of 0.01. The red dashed lines represent a p-value threshold of 0.05. The structures of compounds which enriched with a p-value below 0.01 in both screens are highlighted, along with their activity values from resynthesized compounds in plate-based assays, D_max_ for CDO1 degradation and A_max_ for CDO1-VHL recruitment. *Compound 24 was synthesized and tested as two separate diasteromers. The control compound, VH-032, is also highlighted. Colored by: BB1, Red = Acetyl-t-Leu (Compound 13); Blue = Acetyl-Val (Compound 22); Green = t-Leu Methyl Carbamate (Compound 23); Purple = 1H-pyrrolo[2,3-b]pyridine-5-carboxylic acid (Compound 19). h) Correlation plot between significance of enrichment in stringent splitFAST screen (0.3% gate) and A_max_ values in a plate-based version of the splitFAST assay for 88 orthogonally synthesized compounds. No aggregation of replicate library elements or screen runs was done, logistic curve fit applied, r^2^=0.6 (not shown). Coloring scheme is identical to Figure 5g. Top right quadrant of the confusion matrix represent true positives (34), top left false positives (5), bottom right false negative (0) and bottom left true negatives (50).

Finally, to unambiguously establish on-target mechanism through ternary complex formation, we ran a surface plasmon resonance (SPR) experiment with immobilized CRBN-DDB1 complex with constant flow of 20 μM compound and varying amount of IKZF3-GFP protein^18,71^. Compound 12 induced ternary complex formation with a K_d_ of 8.0 μM (Figure 4g), comparable to the reference degrader pomalidomide (5.1 μM)^71^. Taken together, these data suggest that Compound 12 is a bona-fide CRBN dependent glue degrader. To the best of our knowledge, this compound represents the first example of a succinimide based IKZF3-CRBN glue, a motif which was until recently known only as a CRBN binder and warhead for bifunctionals but not glue degraders^27,72^.

### CDO1-GFP/RFP sensor assay and CDO1-VHL splitFAST biochemical assay in droplets

For a second screen in a more prospective setting, we chose VHL mediated CDO1 degradation^16^, and set out to build a cellular sensor assay analogous to the IKZF3-GFP/RFP sensor assay with a CDO1-GFP fusion and an RFP internal standard (Figure 5a). This cell line proved substantially more challenging to produce, with the CDO1-GFP expression being readily lost in many cell types. Additionally, we found that constiutive expression of CDO1 led to low cell viability in a variety of cell lines. Ultimately, a catalytically dead^73^ CDO1^Y157F^-GFP/RFP construct in HEK293T cells without passage was chosen. Despite being significantly dimmer and the majority of droplets being below detection limit in FADS and therefore ending up in the flow-through regardless of the true activity (Figure 5b, lower left), a clear assay window was still observed with higher fluorescence droplets (Figure 5b, top right to lower right) 3 hours post treatment with DMSO vs. 10 μM VH-032. While suboptimal, we decided to proceed with this assay, based on the premise that statistical analysis enabled by screening more library equivalents might overcome the low assay signal, serving as a good test for the sensitivity and robustness of our workflow by representing a worst-case scenario.

To complement the cellular CDO1-GFP/RFP sensor assay, an orthogonal biochemical CDO1-VHL recruitment assay was developed based on a reversible split fluorescent reporter system^74^ (Figure 5c). The two components of the splitFAST assay were prepared as separate proteins, with the N-terminal fragment of splitFAST (NFAST) fused with CDO1 and the C-terminal fragment of splitFAST (CFAST) fused to VHL co-expressed with Elongin B/C complex. The VHL glue VH-032 induces proximity of CDO1 with VHL, promoting complementation of split components to form FAST protein, which in turn activates the exogenous fluorogenic chromophore (fluorogen) to become highly fluorescent upon its binding. The assay optimization was performed in droplets using FADS. The assay showed an excellent window between the DMSO control and VH-032 in droplets (Figure 5d).

### Prospective 18,432-element VHL-biased library with 18,240 unique members

Next, we designed a VHL-biased 18,432 element (96 acids x 192 amines) library based on a *trans*-L-hydroxyproline core (Figure 5e). Unlike the previous PoC library, only 1 acid building block, acetyl *tert*-leucine (Compound 13), which is featured prominently in VHL bifunctional degraders and CDO1 glues^16^, was represented in duplicate (resulting in 95 unique acids). All 192 amine building blocks were unique, yielding 18,240 unique building block combinations (95 x 192). An explict control compound, the known CDO1 glue degrader VH-032, and several closely analogous amine and acid building blocks were intentionally represented in the library. Analogously to the PoC CRBN-biased library, post solid-phase synthesis qPCR was performed, which indicated an average loading of approximately 42 thousand tags per bead.

### Parallel screens for cellular CDO1 degradation and biochemical CDO1-VHL recruitment with 18,432-element VHL-biased glue library

With the assays and the library in hand, a pair of screens were performed with the VHL library. Using the CDO1-GFP/RFP sensor assay, a cellular screen for CDO1 degradation was performed in 4 independent runs (4,000,000 beads per run, 16,000,000 total, resulting in 800 library equivalents being screened). The screen runs were performed analogously to the previous IKZF3 degradation screen, except with a substantially larger library sampling to account for the low signal intensity of the CDO1 degradation assay. The screen for biochemical CDO1-VHL recruitment was performed using the split-FAST recruitment assay with 2 independent runs (500,000 beads per run, 25 library equivalents), one with a more stringent selection gate (0.3%) and one with a less stringent gate (1.5%). All sorted beads from the selection gates were collected and sequenced as well as 100,000 flow-through beads per run. The collected beads from each experiment were processed in a manner analogus to previous IKZF3 degradation screen, followed by a single NGS experiment for all CDO1-VHL screens, providing raw bead count and enrichment data for further analyses.

A major advantage of the MicDrop platform is the ability to perform orthogonal screens in parallel, providing the ability to compare the cellular degradation hitlist with the biochemical recruitment hitlist to remove potential false-positives and boost confidence in the resulting consensus hitlist prior to initiating resource-intensive resynthesis for further validation. This is visualized by a 2D plot (BB1 vs. BB2) of VHL-CDO1 recruiment (top 0.3% gate), representing CDO1 recruiters with a significance of enrichment (p<0.01) with a square and highlighting CDO1 degraders with a significance of enrichment (p<0.01) with a blue square (Figure 5f). We chose the top 0.3% gate for biochemical screen analysis over top 1.5% gate, because it was found to be more informative. From this analysis, the CDO1 degradation screen was able to correctly identify enrichment of the positive control VH-032 in one of the two instances (p = 0.0116, just below color coding cutoff) as a consensus hit significantly enriched in both screens, along with its fluorinated analogs Compound 14 and 15, as well as three additional consensus hits that were highly enriched in one of the two instances (p <0.01, Compounds 16-18). Interestingly, an unexpected azaindole BB1 (Compound 19) gave two consensus hits that were highly enriched (p <0.01), in addition to two singleton hits with previously unknown motifs (p <0.01), warranting further follow-up (Figure 5f).

When analyzing the screen for biochemical CDO1-VHL recruitment alone (Figure 5f), the building block level analysis revealed an assortment of BB2 combinations being significantly enriched for the two instances of positive control BB1, acetyl *tert*-leucine (Compound 13), and its two analogs, acetyl-valine (Compound 22) and *tert*-leucine methyl carbamate (Compound 23) (Figure 5f). This result is consistent with the known CDO1-VHL recruitment SAR data, although the majority of the exact combinations were previously unknown. In addition, the positive control compound VH-032 showed highly significant enrichment in both replicate library elements (p = 1.42×10^-28^ and 2.30×10^-29^). Furthermore, significant enrichment was largely reproducible between the two instances of acetyl *tert-*leucine, with 31 BB2 combinations being significant (p<0.01) for both cases and only 7 combinations being significant (p<0.01) for only one of the two replicate library elements, confirming the robustness of the biochemical recruitment screen (Figure 5f).

Next, to examine the full correlation of the two screens performed and to determine which compounds to resynthesize, we plotted the significance of enrichment for the cellular degradation vs. the biochemical recruitment screen without library element replicate aggregation (Figure 5g). The consensus hitlist with significant enrichment (p <0.01) for both screens contained 9 compounds (Figure 5g, top right quadrant by green dotted line), with reference compound VH-032 being just below the threshold (p = 0.116) for CDO1 degradation, while highly significant for CDO1 recruitment (p = 1.42×10^-28^ and 2.30×10^-29^). A substantial number of previously unknown building block combinations were enriched significantly only in the CDO1 degradation screen, suggesting that these are off-mechanism false positives likely arising from fluorescence artefacts competing with CDO1-GFP/RFP signals, which were much lower (∼1/10) compared to IKZF3-GFP/RFP system (Figure 5b vs 4b). Thus, even in this non-ideal cellular assay scenario, the orthogonal assay format combination was likely able to rule out many false positives without requiring explicit compound resynthesis and still prioritize bona-fide cell active hits.

### Validation of CDO1-VHL biochemical recruitment and CDO1 cellular degradation screen results

To validate the screen results, we next tested independently synthesized compounds in plate-based CDO1 degradation and CDO1-VHL recruitment assays. First, we examined the 7 consensus hits that were significantly enriched (p <0.01) in both screen formats (Figure 5g). Among those, all 5 compounds bearing acetyl *tert*-leucine BB1 (Compound 13) validated in plate-based versions of the CDO1-VHL recruitment assay and CDO1 degradation assay (Compounds 14-18). By contrast, neither of the other 2 consensus hits with novel BB1 (Compounds 21, 24) showed meaningful activity in either assay formats. The lack of cellular activity of Compound 21 was particularly notable as there are many more azaindole BB1 bearing compounds that were significantly enriched in cellular but not biochemical screens (Figure 4g). Indeed, Compound 20, the top hit from the cellular screen (p = 1.86×10^-8^) but not recruitment screen (p = 1.00), did not show CDO1 degradation in plate format. These data suggest that the CDO1 degradation screen found many false positives, mostly but not exclusively, attributable to the azaindole BB1, while the use of enrichment in biochemical screen as an on-mechanism qualifier likely greatly improved the hit confirmation rate.

We next examined the overall performance of the biochemical assay with all 88 independently synthesized compounds. Compounds 25 and Compound 26, the top biochemical hits from previously underexplored BB1s acetyl-valine (Compound 22) and *tert*-leucine methyl carbamate (Compound 23), respectively, both validated in plate-based format, supporting the robustness of the biochemical screen. The correlation between significance of enrichment in the droplet screen and A_max_ in the plate-based validation was determined by fitting a logistic regression curve (Figure 5h, r^2^ = 0.60, curve not shown). Analogously to the IKZF3 degradation screen, we determined precision and recall using a confusion matrix, with a threshold value for significance of enrichment at a p value of 0.01 in the droplet screen, and a threshold for a true positive at an A_max_ of 500. In this analysis, the precision was 34/39 (87%), with two of the false positives just missing the A_max_ cutoff, while the recall was 100% (34/34). Overall, the biochemical splitFAST screen showed robust performance, with limited confidence in relative activity, but high confidence in the precision and recall, suggesting that we can obtain direct-from-screen SAR from the enrichment data.

### Machine learning activity model based on CDO1-VHL recruitment significance of enrichment and direct-from-screen SAR

Encouraged by the performance of CDO1-VHL recruitment screen that provided direct-from-screen SAR data on 18,240 compounds, we next sought to build a machine learning activity model using Chemprop, a directed message-passing deep neural network model^75^. Chemprop was attractive as an approach, as we felt that the learned encoding of features in the molecular graph representation of structure would be a particularly good fit with a deep data set of compounds in a focused area of chemical space, all sharing a central structural motif. Here, a binary classification model was trained with significance of enrichment data from every library member in the more stringent version of the droplet screen for CDO1-VHL recruitment (0.3% gate) (Figure 6a). We approached model evaluation simultaneously from two perspectives: 1) We sought to employ it as we would in a completely prospective setting, where no SAR is known beforehand and 2) to comprehensively understand model performance, particularly probing our insight into negative data. To satisfy both aims, we predicted the activities of approximately 1,100 readily available compounds containing a hydroxyproline substructure, but critically not present in the training set, and selected the top and bottom 50 compounds for further validation (Figure 6a, bottom right). Finally, the selected 100 independently synthesized compounds were tested in the well-based CDO1-VHL splitFAST recruitment assay and the resulting experimental recruitment A_max_ was plotted against ML model predicted activity (Figure 6b). It is important to note that while the model is trained on significance of enrichment data from a droplet screen, it is being evaluated on its ability to predict the activity of independently synthesized compounds in a well-based version of the same assay.

**Figure 6:**
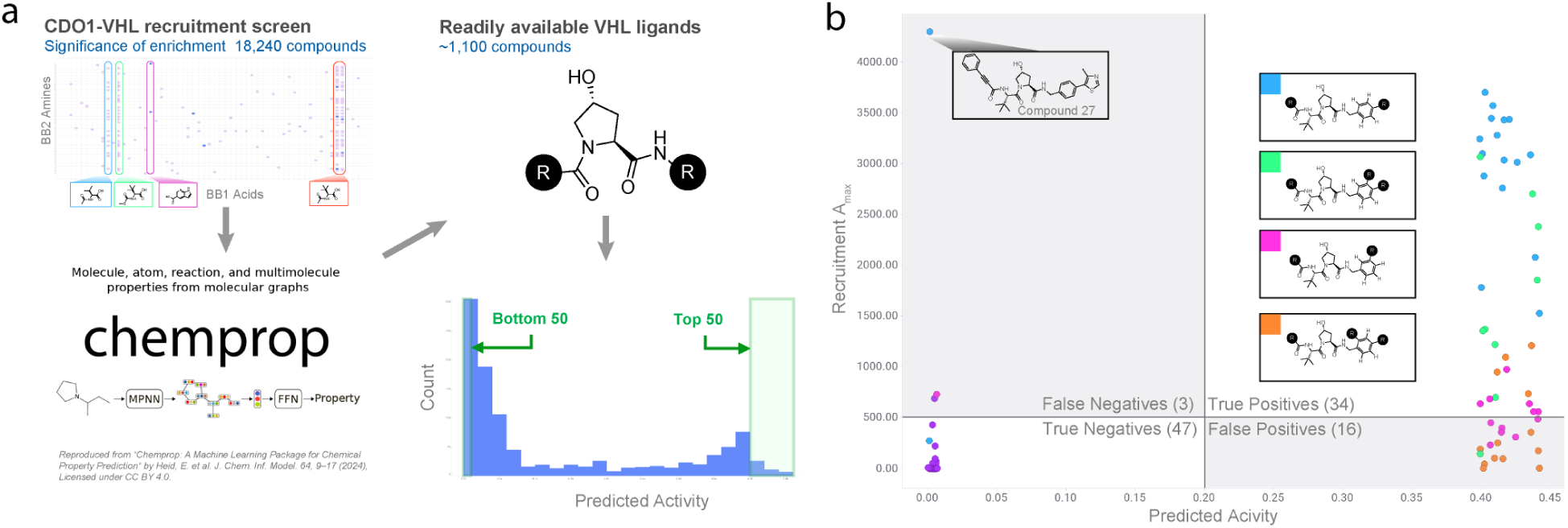
Machine learning model based on CDO1-VHL recruitment direct-from-screen SAR. a) Data from the CDO1-VHL recruitment screen (0.3% gate) was used to train a chemprop classification model, which was benchmarked with only the data from the recruitment screen training set (10 x random split). The model was then applied to ∼1,100 readily available VHL ligands, scoring them on the probability of being classified as active. To test the model, the top 50 and bottom 50 scoring compounds were selected. b) Top 50 and bottom 50 scoring compounds were tested in a plate-based version of the CDO1-VHL splitFAST recruitment assay. A_max_ is plotted against predicted acitivity. Model showed 92% precision and 68% recall. Points on plot are colored by substructure present as described in figure. Chemprop banner (left bottom) reproduced from *“Chemprop: A Machine Learning Package for Chemical Property Prediction“* by Heid, E. et al. J. Chem. Inf. Model. 64, 9–17 (2024), Licensed under <u>CC BY 4.0.</u>

As selecting the top 50 compounds reflects a plausible strategy for utilizing the model in a purely prospective manner, its performance there may be most critical. Gratifyingly, of these top 50 predicted compounds, 34 were found to be CDO1-VHL recruiters, reflecting a precision of 68%. The fact that the majority of compounds were active is encouraging from the perspective of *in-silico* SAR expansion. Predicting the activity of the bottom 50 compounds turned out to be even more accurate, with 47 of the 50 being classified correctly as inactive. The only prominent false negative was Compound 27, a potent recruiter which likely found a narrow exit vector from ternary complex^16^ with a motif absent from the training set library. Overall, the ML model was able to predict recruitment activity with 92% precision and 68% recall, strongly supporting that significance of enrichment from the droplet screen is sufficient to provide direct-from-screen SAR.

Perhaps most intriguingly, the correctly predicted active compounds contained a range of substitution patterns on the benzylamine right-hand side of the molecule. The training set 18,240-member library did not contain fully elaborated representations of these benzylamine substitution patterns, especially bulky groups such as 3,4– and 2,4-disubstitution with bulky aryl groups. The fact that the model correctly predicted that these would be active is in itself interesting, but more compelling is that this represents productive SAR exploration as an additional, unexpected benefit. Taken together, the findings from training this ML model further support that the combination of droplet-based screen and machine learning model can work in concert to rapidly create experimental and virtual SAR.

## Discussion

To the best of our knowledge, this work represents the first demonstration of cellular phenotypic screening of a DNA-encoded combinatorial library in droplets. In contrast to previously reported eOBOC cellular screening approaches utilizing arrayed compartments such as agar lawn^47^, nanopens^49^ or nanowells^15^, we generate millions of water-in-oil microdroplets in a matter of minutes, and pool and sort them continuously in flow. This scalability combined with NGS allows for screening of many bead replicates to enable statistical analysis on contrast of counts. The examples described achieved not only identification of select hits, but also direct-from-screen SAR, including ranking of relative potency in the case of the cellular IKZF3 degradation screen. This is a significant departure from affinity-selection methods^38–40^, which typically lack biological replicates and disregard non-hits. Indeed, the replicate-based, large, high-quality data generated from our workflow was sufficient to directly build and benchmark an actionable ML activity model. It is also worth noting that the campaigns described were the very first that we performed, and many process improvements are conceivable and underway.

In encoded bead library screens, the sampling of individual library members is governed by a normal distribution^63^. We therefore chose to screen the number of library equivalents that we deemed appropriate as an average replicate count per compound^63^. For the cellular IKZF3 degradation screen, 100 library equivalents were screened (before replicate library element aggregation). The fact that we observed substantially better correlation with well-based assay data after replicate library element aggregation highlights the value of greater bead replicates and redundant syntheses in our workflow. This is similar to contrast of counts-based screening technologies such as pooled CRISPR screens, where average sampling per guide RNA per sample routinely ranges 100-1,000 to detect subtle, but statistically significant candidate hits^76^. Our multiple library element replication (synthesis replicates, each of which are encoded with a unique compound specific barcode, CSB) could also mirror pooled CRISPR screens where each gene is tested with independent guide RNAs. In addition, because of a challenging assay for the CDO1 degradation screen, we screened 800 library equivalents to maximize our statistical power, whereas for the biochemical CDO1-VHL recruitment assay with a large assay window, we screened only 25 library equivalents and achieved robust statistics.

While we set out to perform PoC screens expecting to rediscover known compounds and close analogs in the IKZF3 degradation screen, the unexpected discovery of a singleton hit, Compound 12, from the succinimide motif included *explicitly intending to produce only inactive combinations* highlights the value of bespoke combinatorial libraries, where the assumed low probability of success lies beyond the scope of a conventional library build paradigm, but within the purview of full matrix enumeration. This compound was not present in our traditional chemistry-based 50K CRBN library at Novartis and there were no succinimide-based IKZF3 glue degraders known internally at Novartis. The unexpected finding is particularly encouraging considering the limited diversity of a set of only 364 compounds, suggesting that the prevalence of such findings when screening much larger, more diverse libraries should be significantly greater.

It is worth noting that the MicDrop platform built for cellular functional assays was readily applicable to biochemical assays as demonstrated by the splitFAST CDO1-VHL screen. The lack of cell-to-cell and droplet-to-droplet fluorescence variability enabled us to screen fewer library equivalents to achieve excellent screen statistics, precision and recall. Interestingly, however, the biochemical recruitment screen showed a more binary distribution of hits and non-hits and a reduced ability to rank order relative potency of the reference compound when compared with the cellular IKZF3 degradation screen (Figure 5h vs. Figure 4e). This may be due to the larger assay window and tighter response to a given compound that produces a binary rather than continuous distribution, which may be addressed by introducing elements of dose response, such as protein concentration, multiple selection gates, and varying compound loading or UV release^58,77^. Another explanation for the observed binary hit distribution may be a possible avidity effect from the splitFAST tags.

Massive miniaturization of both chemistry and screening empowers our platform to perform parallel screens with orthogonal assays, increasing confidence in hits while progressing further down the drug optimization flowchart without the need for costly and time-consuming compound resynthesis. This was particularly valuable for the CDO1 degradation screen, where the vast majority of likely false positives arising from low assay signal could be filtered out by simply applying a minimum threshold value for biochemical recruitment (Figure 5f,g). For protein degradation screens with a CRBN-biased library, additional parallel screens could include off-target profiling via GSPT1^7^ and SALL4^13^ degradation and on-mechanism validation with CRBN knock-out or NAEi treatment^17^. Besides boosting confidence in the obtained hits, direct-from-screen SAR knowledge generated for counter targets should further facilitate subsequent optimization efforts, especially by building predictive off-target ML models to avoid synthesis of analogs with undesired off-target profiles.

Some of the pending improvements to the platform described in this manuscript include: increasing the throughput of the droplet sorter, which determines the screenable library size, additional assay readouts beyond two-color fluorescence, hydrogel matrix for adherent cells and also enabling nutrient exchange and addition of detection reagents post droplet breakage while keeping phenotype and genotype intact. For the throughput, the current setup with a 500Hz FADS allows routine screens of up to 10 million beads per week, which translates to cellular degradation screens of 100K member libraries with 100 library equivalents, or biochemical recruitment screens of 400K member libraries with 25 library equivalents.

Further improvement in FADS, parallelization and optimization of droplet workflow provide possible avenues for increasing throughput. For broader scope of cells and assays, adoption of hydrogel particles or core-shell beads^78^ may be one avenue to improve throughput while allowing reagent addition and compatibility with a broader set of instruments, such as wide-nozzle flow cytometers and cell pickers with various imaging readouts.

While recent advances in artificial intelligence driven by sophisticated deep learning models are breathtaking, there is an increasing realization of data scarcity, that large and curated data sets in the relevant domain are urgently needed^1^. To this end, our cellular and biochemical functional screens driven by high replicate counts provide large direct-from-screen SAR data which can directly train target-specific ML activity models. The potential implication is quite significant, considering the traditional bottleneck of drug discovery has been the synthesis and testing of discrete compounds to slowly build SAR over the course of years with several hundreds to thousands of compounds. Combining our workflows for empirical screening of 10s to 100s of thousands of compounds with machine learning models could enable multiple, semi-virtual, design-make-test-analyze (DMTA) cycles prior to more resource intensive, traditional lead optimization towards a clinical candidate.

In summary, we set forth a platform that enables scalable cellular and biochemical functional screens of large combinatorial libraries in droplets. While MicDrop was developed to accelerate the discovery of molecular glues for chemical induced proximity modalities, it has broader implications for hit finding and optimization, suggesting a possible shift in the screening paradigm from hit finding to lead discovery with actionable SAR, accelerating the path to development candidates and substantially reducing time and resource requirements. We also believe this work provides a new framework for further optimization and standardization to enable broader adoption of droplet-based screens of combinatorial libraries. Finally, the ability to interrogate cellular function with large, fit-for-purpose bespoke libraries has the potential to transform the traditional approach of screening what is available to what is imaginable, providing access to the vast chemical whitespace, democratizing drug discovery and unlocking elusive drug targets to deliver novel therapeutics.

## Acknowledgements

The authors thank colleagues from On-Chip Biotechnologies for optimizing On-Chip Sort for our droplet screen workflow: Mr. N. Honma and Mr. K. Akita for on-site support; Mr. K. Ishikawa, Dr. Y. Fujimura and the entire engineering team for the development of the continuous feed adaptor. The authors also thank Professor B. Paegel for his pioneering eOBOC work and early consultancy sessions to orient us with the basics of eOBOC library technology. The authors also thank many Novartis colleagues whose efforts enabled this work: D. Vickers, K. Choquette, D. Evans, A. Meyer, J. Mainquist and D. Sipes for prototyping the first generation droplet sorter; W. Partlo and K. Scott for building the second generation sorter and performing benchmarking studies with an alternative assay; D. Casalena for prototyping biochemical assays in droplets; N. Blanks and T. Burks for hardware support; D. Nunes and A. Ho for performing flow cytometry experiments; V. Ruda, L. Mansur, T. Burks, C. Schwalen, J. Rigal, S. Baker for their support on PCR and NGS experiments; B. Leon and D. Dovala for protein production support; J. McKenna M. Eberle and R. Tichkule for CRBN building blocks and library support; M. Zambrowski for mass spectroscopy support; A. Schuffenhauer and L. Shen for chemoinformatic support; C. Dickson for CADD support; W. Ulmer for building block dissolution and QC; M. Fiorino for custom plate-based synthesis setup; A. Bickel and C. Petersen for the UV dosing chamber design support; D. Buckley and G. Michaud for VHL-CDO1 SAR insights and building blocks; A. Abrams for figure artwork support; C. Lee for initial insights and continued support; G. Hoffman for gathering initial support; F. Berst for helpful discussions and support; Y. Isome for project management; N. Dales and T. Gabriel for early project mentorship; TPD devotees W. Forrester, G. Hollingworth, J. Solomon and R. Beckwith for countless discussions on accelerating molecular glue discovery; former and current Novartis leadership: J. Bradner, K. Briner, H. Hemmerle, R. Jain, J. Quancard, S. Canham, M. Mogi and J. Tallarico for their encouragement and support, J. Joslin and J. Ottl for their critical feedback and support; Genesis Labs: I. Hunt, A. Reynolds, M. Chaffers and A. Spier who together provided the initial sandbox for MicDrop as an internal incubator project; and C. Dumelin, G. Hollingworth, C. Lee, J. Solomon and P. Ting for careful review of the manuscript.

## Funding

This study was supported by Novartis Biomedical Research.

## Author Contributions

M.T., L.W., F.D.S., G.E., A.A.A. and K.Y. designed and performed end-to-end workflows as described. C.F., A.A.A., H.I., L.W. and K.Y. prototyped and M.T. optimized bead chemistry. C.F. designed and C.F. and F.D.S. validated eOBOC encoding scheme. C.F. and K.Y. designed and C.F. validated and prepared prototype CRBN library. M.T. and K.Y. designed, M.T. and L.W. validated and prepared CRBN and VHL eOBOC libraries for the studies described. G.E. developed custom microfluidic setup for droplet generation and UV dosing, including PDMS chip design and production capabilities. A.A.A. developed emulsion formulation for biochemical assays. A.A.A., B.M., T.A. and K.Y. prototyped microfluidic bead encapsulation and M.T., L.W., G.E. and K.Y. optimized co-encapsulation of beads and cells into droplets. A.A.A., H.I., F.M., K.Y. prototyped and M.T., L.W., G.E. optimized UV-linker and release conditions. G.E., P.S. prototyped, and A.B. optimized custom UV dosing devices. S.C. created and S.P. developed IKZF1-GFP/RFP assay in Jurkat cells. S.P. validated in-drop IKZF3 degradation with post-breakage FACS analysis. D.S.A. prototyped splitFAST assay for detecting ternary complex formation. A.T. developed VHL-CDO1 splitFAST and CDO1-GFP/RFP assays. A.A.A. optimized CDO1-VHL splitFAST assay in droplets. B.M., A.A.A. and A.K.P. prototyped in-drop IKZF3 degradation by microscopy where A.K.P. designed and optimized the imaging protocol and image analysis. M.T. and G.E. performed in-drop IKZF3 degradation studies by imaging as described. S.P. prototyped OnChip Sort as off-the-shelf FADS. M.T. optimized the workflow and gating strategy using OnChip Sort for optimal screening performance. M.T. and L.W. performed IKZF3 and CDO1 degradation as well as CDO1-VHL recruitment screens using OnChip Sort. C.F. developed and L.W. optimized eOBOC PCR protocols. L.W. prepared samples and FDS designed and supervised NGS experiments by external provider. F.D.S. devised NGS data pipeline, performed statistics & hit calling, and visualized data for all screens described in this manuscript. F.D.S., M.T. and K.Y. analyzed the screen data and designed screen result validation studies including droplet to microtiter well plate-based assay correlation analysis. M.T. performed hit re-synthesis for all screens. D.S.A. and G.E. performed microtiter well plate-based hit validation assays for IKZF3 degradation screen. S.C. designed and J.H. performed capillary Western experiment for IKZF1, IKZF3 and GSPT1 degradation +/− NAEi, and C.A.W. performed SPR assay to characterize novel succinimide-based IKZF1 hit Compound 12. A.T. and L.A. performed microtiter well plate-based hit validation assays for CDO1 degradation and recruitment. M.T. and J.F. designed and J.F. performed the ML experiment. A.T. and L.A. performed microtiter well plate-based ML hit validation. K.Y. conceived the MicDrop project and oversaw the work described. M.T. and K.Y. wrote this manuscript.

## Competing interests

All authors are past or current employees of Novartis. Some of the authors have patents related to this work: WO22190014 A1, High-throughput screen in droplets (A.A.A., T.A., G.E., B.M., S.P., and K.Y.). The authors declare no other competing interests.

